# ScRNAbox: Empowering Single-Cell RNA Sequencing on High Performance Computing Systems

**DOI:** 10.1101/2023.11.13.566851

**Authors:** R.A. Thomas, M.R. Fiorini, S. Amiri, E.A. Fon, S.M.K. Farhan

## Abstract

**Motivation:** Single-cell RNA sequencing (scRNAseq) offers powerful insights, but the surge in sample sizes demands more computational power than local workstations can provide. Consequently, high-performance computing (HPC) systems have become imperative. Existing web apps designed to analyze scRNAseq data lack scalability and integration capabilities, while analysis packages demand coding expertise, hindering accessibility.

**Results:** In response, we introduce scRNAbox, an innovative scRNAseq analysis pipeline meticulously crafted for HPC systems. This end-to-end solution, executed via the SLURM workload manager, efficiently processes raw data from standard and Hashtag samples. It incorporates quality control filtering, sample integration, clustering, cluster annotation tools, and facilitates cell type-specific differential gene expression analysis between two groups.

**Implementation:** Open-source code and comprehensive usage instructions with examples are available at https://neurobioinfo.github.io/scrnabox/site/.

**Supplementary Information:** Supplementary data are available at Bioinformatics online.

## Introduction

In recent years, single-cell RNA sequencing (scRNAseq) technology has led to remarkable breakthroughs in our understanding of biology, enabling us to explore gene expression at the resolution of individual cells. With technological advancements, we have transitioned from analyzing a few cells to thousands and even hundreds of thousands of cells in a single experiment (1). While the potential of scRNAseq is immense, it has brought about complexities and computational demands that have yet to be comprehensively addressed. Many useful web-based applications and graphical user interfaces (GUI) have been developed to analyze scRNAseq; however, with the exception of Asc-Seurat, these tools require processing of each sample separately, therefore, inhibiting data integration or comparisons among experimental variables (2–6). R and python packages with excellent user guides have been developed to process scRNAseq data; however, these require extensive programming knowledge (7–11). Additionally, given that users must manually adapt and implement the code, repeating the process for each sample, this process can be laborious, error-prone, and time-consuming. The need to execute the code locally further exacerbates these issues, limiting researchers to the capabilities of their own computational resources. The scale of modern scRNAseq datasets necessitates the use of high-performance computing (HPC) clusters. Yet, to our knowledge, a comprehensive scRNAseq workflow tailored to HPC environments hitherto has been unavailable.

In response to these multifaceted challenges, we introduce scRNAbox, a novel and robust scRNAseq analysis pipeline meticulously designed for HPC systems. ScRNAbox not only standardizes and simplifies the scRNAseq analysis workflow for geneticists and biologists with any levels of computational expertise, but also diligently documents execution parameters, ensuring transparency and replicability. It has been assembled to be effortlessly scalable, catering to the evolving needs of researchers faced with large-scale datasets. scRNAbox provides a unified and accessible resource for the growing community of scRNAseq researchers.

Finally, we recognize the shortage of resources that provide best practices in scRNAseq analysis (12, 13). In this context, we deploy scRNAbox using publicly available data and outline the decisions bioinformaticians must make during analysis and investigate the biology. To illustrate the utility of scRNAbox, we analyze single-nuclei RNA sequencing (snRNAseq) data published by Smajic and colleagues of midbrain tissue from patients with Parkinson’s Disease (PD) and controls (14). We outline each step in the scRNAbox pipeline, providing the scientific rationale and the analytical decisions taken in processing the data.

## Materials and methods

### scRNAbox overview

The scRNAbox pipeline consists of R scripts that are submitted to the SLURM workload manager (job scheduling system for Linux HPC clusters) using bash scripts from the command line. (15). Beginning with 10X Genomics expression data from raw sequencing files, the pipeline facilitates standard steps in scRNAseq processing through to differential gene expression between two different conditions. The scRNAbox framework consists of three main components: (i) R scripts, (ii) job submission scripts, and (iii) parameter and configuration files. The pipeline is separated into Steps, which correspond to analytical tasks in the scRNAseq analysis workflow (**Figure 1**). Users can tailor their analysis by manipulating the parameters in the step-specific parameter files. The pipeline can analyze scRNAseq experiments where each sample is captured separately (standard track) or multiplexed experiments where samples are tagged with sample-specific oligonucleotide tagged Hashtag antibodies (HTO), pooled, and sequenced together (HTO track) (16, 17). The results of each step are reported in intuitive tables, figures, and intermediate Seurat objects (7). Upon submitting the bash script for a step, “Jobs”, or resource requests are created based on the parameters defined in the configuration file, including CPUs, memory, and time. Jobs are submitted to the HPC system using the SLURM “Scheduler” to execute the R scripts. A complete user guide and the code used in this manuscript can be found at the scRNAbox GitHub site: https://neurobioinfo.github.io/scrnabox/site/.

**Figure 1.**
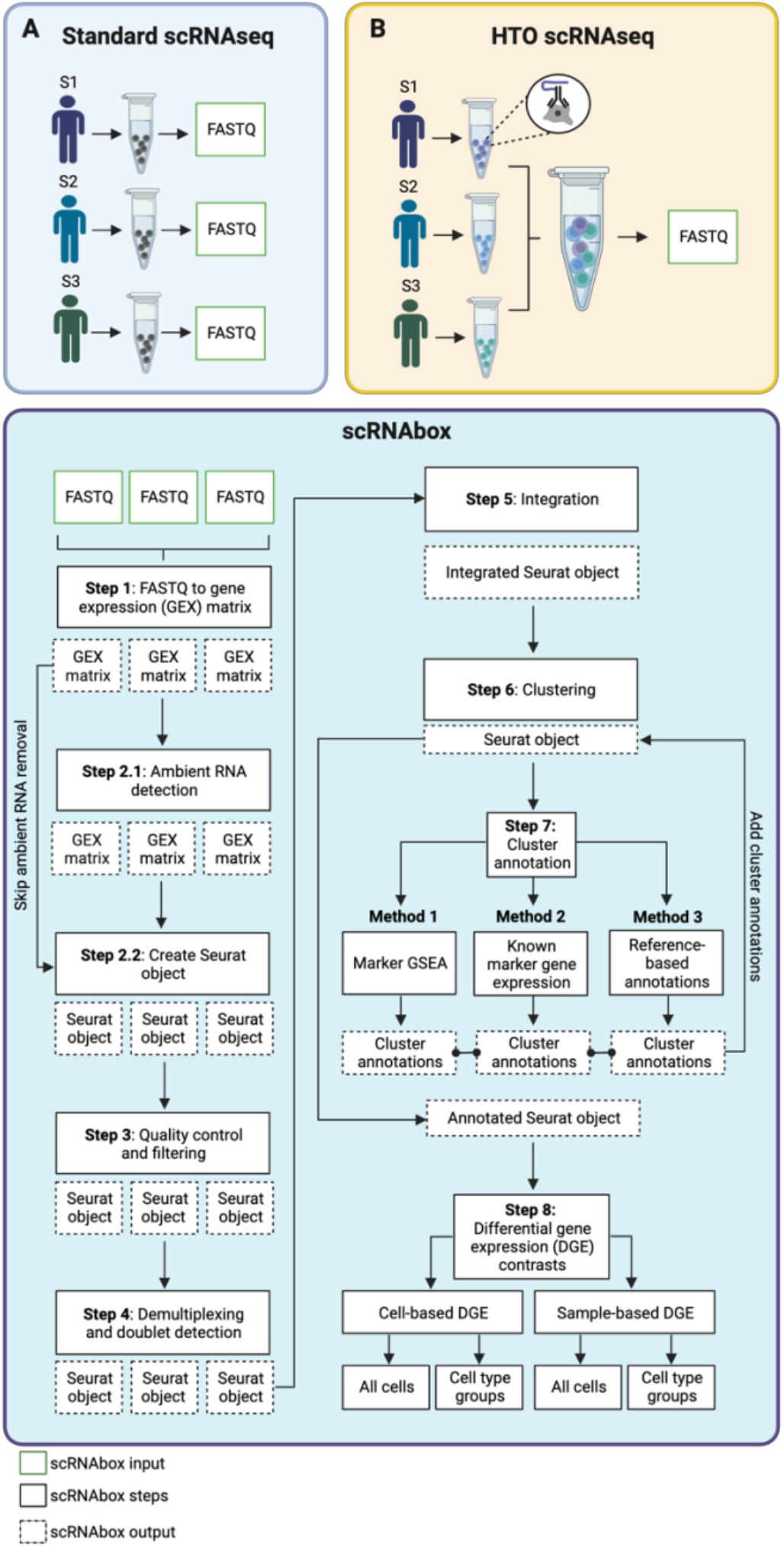
ScRNAbox analysis workflow. The scRNAbox pipeline provides two analysis tracks: 1) standard sRNAseq and 2) HTO scRNAseq. **A)** Standard scRNAseq data is prepared by sequencing each sample separately, resulting in distinct FASTQ files for each sample. **B)** HTO scRNAseq data is produced by tagging the cells from each sample with unique oligonucleotide “Hashtag” conjugated antibodies (HTO). Tagged cells from each sample are then pooled and sequenced together to produce a single FASTQ file. Sample-specific HTOs are used to computationally demultiplex samples downstream. **C)** Steps of the scRNAbox pipeline workflow: steps are designed to run sequentially and are submitted using the provided bash scripts through the command line. scRNAbox takes FASTQ files as input into Step 1; however, the pipeline can be initiated at any step which takes the users processed data as input.

### Installation

ScRNAbox can be installed on any HPC Linux system via the *scrnabox.slurm* package, which contains the Bash and R scripts, parameter files, and configuration file. The HPC system must have CellRanger (<u>10xGenomics</u>) and R (v4.2 or higher) (18) installed and must use a SLURM scheduler. Users must also install several R libraries using the provided installation code.

### scRNAbox Steps

#### Step 0: Initiation and configuration

Following installation, users run Step 0 to initiate the pipeline and specify if they will use the standard or HTO analysis track. Step 0 creates the job submission configuration files and the step-specific parameter files. The configuration file contains the time and memory usage settings for each step and must be edited to match the user’s needs. After Step 0, each subsequent Step can be run individually through separate commands or all together in a single command.

#### Step 1: FASTQ to gene expression matrix

##### 1.1 : File structure and inputs

Prior to running the CellRanger *counts* pipeline, a parent directory (“samples_info”) must be created in the working directory. The “samples_info” directory must contain a folder for each sample; the name of the sample-specific folders will eventually be used to name the samples in downstream steps. Each sample-specific folder must contain a *library.csv* file, which defines the information of the FASTQ files for the specific sample. The HTO analysis track also requires a *feature_ref.csv* file, which specifies the oligonucleotide sequences of the Hashtags. Step 1 runs a script to automatically generate these files based on the user input in the parameter file. However, users can manually generate the required files and structure.

##### 1.2 : Running CellRanger

ScRNAbox deploys the CellRanger *counts* pipeline to perform alignment, filtering, barcode, and unique molecular identifier counting on the FASTQ files. Each sample is processed by the CellRanger *counts* pipeline in parallel. Although CellRanger is processed with default parameters, all relevant parameters can be adjusted (<u>10X Genomics</u>).

#### Step 2: Create Seurat object and remove ambient RNA

##### 2.1 : Ambient RNA detection

The R package SoupX is used to account for ambient RNA, providing users the option to correct the gene expression matrices for RNA contamination (19). SoupX quantifies the contamination fraction according to the expression profiles of empty droplets and cell clusters identified by the CellRanger *counts* pipeline. Marker genes used to estimate the contamination rate are automatically identified using the *AutoEstCont* function and the expression matrix is corrected per the estimated contamination rate using the *adjustCounts* function.

##### 2.2 : Generation of the Seurat object and quality control metrics

The Seurat function *CreateSeuratObject* is used to take in the CellRanger (if not removing ambient RNA) or SoupX (if removing ambient RNA) generated feature-barcode expression matrices, and create the list-type Seurat object (7). The number of genes expressed per cell (number of unique RNA transcripts) and the total number of RNA transcripts are automatically computed. The proportion of RNA transcripts from mitochondrial DNA (gene symbols beginning with “MT”) and the proportion of ribosomal protein-related transcripts (gene symbols beginning with “RP”) are both calculated using the Seurat *PercentageFeatureSet* function. Following the Seurat workflow, the *CellCycleScoring* function with the Seurat S and G2/M cell cycle phase reference genes are used to calculate the cell cycle phase scores and generate a principal component analysis (PCA) plot (20).

#### Step 3: Quality control and generation of filtered data objects

ScRNAbox allows users to filter low quality cells by defining upper- and lower-bound thresholds in the parameter files based on unique transcripts, total transcripts, percentage of mitochondrial-encoded transcripts, and percentage of ribosome gene transcripts. Users can also remove or regress a custom gene list from the dataset. The filtered counts matrix is then normalized, the top variably expressed genes are identified, and the data are scaled using Seurat functions. Linear dimensional reduction is performed via PCA and an elbow plot is generated to visualize the dimensionality of the dataset and inform the number of principal components (PC) to be used for doublet detection in Step 4.

#### Step 4: Demultiplexing and doublet removal

##### 4.1 Doublet detection and removal (Standard track)

Barcodes that are composed of two or more cells are identified as doublets using DoubletFinder (21). Doublets are predicted based on the proximity of each cell’s gene expression profile to that of artificial doublets created by averaging the transcriptional profiles of randomly chosen cell pairs. The default value of 0.25 for the number of artificial doublets is used. The neighbourhood size corresponding to the maximum bimodality coefficient is selected and the proportion of homotypic doublets is computed using the *modelHomotypic* function. Users can define the number of PCs to use for doublet detection and the expected doublet rate for each sample. Users have the option to remove doublets from downstream analyses or just calculate the doublet rate.

##### 4.2 Demultiplexing followed by doublet removal (HTO track)

Pooled samples are demultiplexed, assigning an HTO label to each cell, using Multi-seq (17). The automatically detected inter-maxima quantile thresholds of the probability density functions for each barcode are used to classify cells. Cells surpassing one HTO threshold are classified as singlets; cells surpassing >1 thresholds are classified as doublets; the remaining cells are assigned as “negative”. The counts observed for each barcode are reported in a summary file and plots are generated to visualize the enrichment of barcode labels across sample assignments. Users have the option to remove doublets and negatives from downstream analyses.

#### Step 5: Creation of a single Seurat object from all samples

##### 5.1 : Integration or merging samples

The individual Seurat objects are integrated to enable the joint analysis across sequencing runs or samples by deploying Seurat’s integration algorithm (22). The genes that are variable across all samples are detected by the *SelectIntegrationFeatures* function. Integration anchors (pairs of cells in a matched biological state across datasets) are selected by the *FindIntegrationanchors* function, and the *IntegrateData* function is used to integrate the datasets by taking the integration anchors as input. Alternatively, users may simply merge the normalized counts matrices using Seurat’s *merge* function without performing integration.

##### 5.2 : Linear dimensional reduction

Seurat functions are used to normalize the count matrix, find the most variably expressed genes, and scale the data. Linear dimensional reduction is then performed via PCA using the top variably expressed genes as input. An elbow plot to visualize the variance contained within each PC and jackstraw plot to visualize “significant” PCs are produced. These plots inform the number of PCs that should be retained for clustering in Step 6.

#### Step 6: Clustering

Clustering is performed to define groups of cells with similar expression profiles using the Seurat implementation of the Louvain network detection with PCA dimensionality reduction as input (7). K-nearest neighbours are calculated and used to construct the shared nearest neighbour graph. The Jaccard similarity metric is used to adjust edge weights between pairs of cells, and the Louvain algorithm is used to iteratively group cells together based on the modularity optimization. To assist users in selecting the optimal clustering conditions, we include an option to compute the Louvain clustering N times at each clustering resolution, while shuffling the order of the nodes in the graph for each iteration. The average and standard deviation of the Adjusted Rand Index (ARI) between clustering pairs at each clustering resolution is then calculated (23). A *ClustTree* plot (24) and uniform manifold approximation and projection (UMAP) plots are generated to visualize the effect of clustering parameters.

#### Step 7: Cluster annotation

Cluster annotation is performed to define the cell types comprising the clusters identified in Step 6. ScRNAbox provides three tools to identify cell types comprising the clusters.

##### 7.1 : Tool 1: Cluster marker gene identification and gene set enrichment analysis

ScRNAbox identifies genes that are significantly up regulated within each cluster by using the Seurat *FindAllMarkers* function, implementing the Wilcoxon rank-sum test (7), with log2 fold-change (L2FC) threshold of 0.25. Differentially expressed genes (DEGs) are calculated by comparing each cluster against all the other clusters, only upregulated genes are considered. A heatmap is generated to visualize the expression of the top marker genes for each cluster at the cell level. All significant DEGs are used as the input for gene set enrichment analysis (GSEA) across user-defined libraries that define cell types using the EnrichR tool (25). Cluster-specific tables are generated to report all enriched cell types and bar plots visualize the most enriched terms.

##### 7.2 : Tool 2: Expression profiling of cell type markers and module scores

ScRNAseq allows users to visualize the expression of individual genes and the aggregated expression of multiple genes from user-defined cell type marker gene lists. For each gene in a user-defined list, a UMAP plot visualizes its expression at the cell level, while violin and dot plots visualize its expression at the cluster level. Aggregated expression of user-defined cell type marker gene lists is calculated using the Seurat *AddModuleScore* function (20). The average expression of each cell for the gene set is subtracted from randomly selected control genes, resulting in cell-specific expression scores, with larger values indicating higher expression across the gene set.

##### 7.3. Tool 3: Cell type predictions based on reference data

ScRNAbox utilizes the Seurat label transfer method: *FindTransferAnchors* and *TransferData* functions, to predict cell-type annotations from a reference Seurat object (22). Predicted annotations are directly integrated into the query object’s metadata and a UMAP plot is generated to visualize the query dataset, annotated according to the predictions obtained from the reference.

##### 7.4 Adding annotations

ScRNAbox uses the Seurat *AddMetaData* function and a user-defined list of cell types in the parameter file to add cluster annotations. The cluster annotations from each iteration of the step will be retained, allowing users to define broad cell types and subtypes. UMAP plots with the annotation labels are generated to visualize the clustering annotations at the cell level, allowing users to check the accuracy of their annotations.

#### Step 8: Differential gene expression analysis (DGE)

Metadata defining the groups to be compared are added to the Seurat object by submitting a .csv file containing sample information with phenotypic or experimental data. The additional metadata is used to define the variables to compare for the DGE. ScRNAbox allows DGE to be calculated between conditions using all cells or cell type groups using two different data preparations: cell-based or sample-based DGE.

##### 8.1 : Cell-based DGE

Cells are used as replicates and DGE is computed using the Seurat *FindMarkers* function to compare user-defined contrasts for a given variable (7). While *FindMarkers* supports several statistical frameworks to compute DGE, we set the default method in our implementation to MAST, which is tailored for scRNAseq data (26). MAST models both the discrete expression rate of all genes across cells and the conditional continuous expression level, which is dependent on the gene being expressed in the cell, by a two-part generalized linear model (26). Regardless of the method used, P values are corrected for multiple hypothesis testing using the Bonferroni method. Users can perform their own p-value adjustments using the DEG files output from the pipeline.

##### 8.2 : Sample-based DGE

To calculate DGE using samples or subjects as replicates, scRNAbox applies a pseudo-bulk analysis. First, the Seurat *AggregateExpression* function is used to compute the sum of RNA counts for each gene across all cells from a sample (27). These values are then input into the DESeq2 framework, which uses gene dispersal to calculate DGE (28). P values are corrected for multiple hypothesis testing using the Bonferroni method, which can be recalculated from the pipeline output.

##### 8.3 : Analysis of differentially expressed genes

Step 8 produces data tables of the DEGs for each of the defined contrasts. These outputs can be used for gene enrichment pathway analysis using web-apps or though application program interfaces with reference libraries using a programming language, in our case, R. Further analysis of the results is experiment-dependent and must be completely tailored to the research questions. We used the ClusterProfiler R package to identify significantly enriched Gene Ontology (GO) terms with the *gseGO* function (29). We utilized the ‘org.Hs.eg.db’ Bioconductor annotation package to access human (*Homo sapiens*) gene annotations for our analysis. The ggplot2 R package was used for data visualization (30).

### Code and data availability

The *scRNAbox.slurm* package is available at https://github.com/neurobioinfo/scrnabox. All code is open source and licenced under the MIT license, allowing for reuse, alteration, and sharing with credit given. Bash submission scripts, R scripts, configuration files, parameter files, and R scripts used for DGE analysis and to generate all the plots in the figures are available. We analyzed publicly available data summarized in **Table 1**.

**Table 1:** Description of datasets.

| Data type | Source | Reference |
| --- | --- | --- |
| SnRNAseq post-mortem human midbrain | GEO: GSE157783 | Smajic et al. (14). |
| ScRNAseq human PBMC with Hashtags | GEO: GSE108313. | Stoeckius et al. (16). |
| SnRNAseq from the post-mortem human midbrain | https://singlecell.broadinstitute.org | Kamath et al (31). |

Abbreviations: GEO, Gene Expression Omnibus; PBMC, peripheral blood mononuclear cell; scRNAseq, single-cell RNA sequencing; snRNAseq, single-nucleus RNA sequencing.

## Results

To demonstrate the functionality of the scRNAbox pipeline we analyzed a publicly available snRNAseq dataset from the post-mortem midbrains of five patients with Parkinson’s disease (PD) and six controls prepared by Smajic et al. (14). To demonstrate scRNAbox’s ability to process multiplexed scRNAseq data, we analyzed a scRNAseq dataset of peripheral blood mononuclear cells (PBMCs) from eight human donors prepared by Stoeckius et al. (16).

### ScRNAbox efficiently processes raw sequencing data and provides quality control measures

We initiated our scRNAbox analysis of the midbrain dataset by running Step 0 and selecting the standard track. This created the job configuration file and step-specific parameter files. In Step 1, we used the automatic library preparation function to generate the sample-specific *library.csv* files and ran CellRanger (v5.0.1) *counts* on all 11 subjects.

In Step 2, the pipeline generates a Seurat object for each sample and computes multiple quality control metrics that inform decisions for filtering in Step 3. At this stage, we had the option to remove ambient RNA, transcripts from an external source captured with a true cell. These aberrant transcripts originate from many possible sources including cells that ruptured or died during dissociation and released their RNA, mRNA-containing exosomes, or mRNA that leaked out when cell processes were cleaved during dissociation. Large amounts of ambient RNA confound the data, making cells appear to have similar transcriptional profiles when they are truly distinct. Leveraging SoupX to detect ambient RNA revealed low contamination rates across all samples (mean = 2.46%) (**Figure 2A**; **Table 2**). Cell cycle stage is another quality control metric to consider during scRNAseq data processing as it can affect cell type annotations in downstream analyses. ScRNAbox computes the cell cycle stage for each cell and generates a PCA plot to visualize the effect of cell cycle stage in the data. The cell cycle stage showed little effect on cell distributions in PCA space (**Figure 2B; Supplementary Figure S1**).

**Figure 2.**
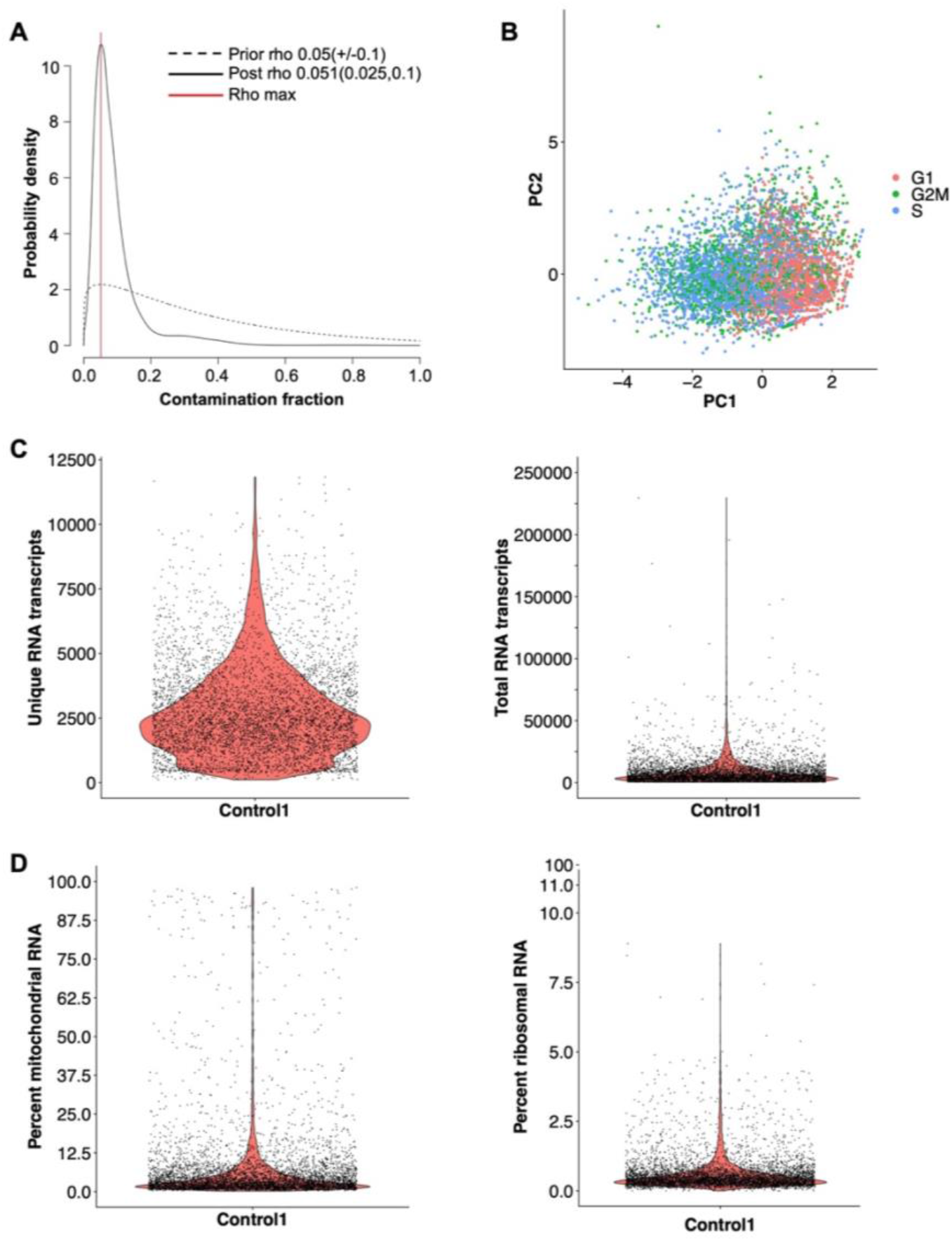
scRNAbox calculates and visualizes quality control metrics. **A)** Line plot of the ambient RNA contamination rate (rho) estimated by SoupX (19). Estimates of the RNA contamination rate using various estimators are visualized via a frequency distribution; the true contamination rate is assigned as the most frequent estimate. The ambient RNA rate for snRNAseq midbrain sample *Control 1* is indicated by the red line (5.1 %). **B**) Scatter plot of PCA analysis of *Control 1* coloured by cell-cycle scores calculated using the Seurat S and G2/M reference genes (20). **C)** Violin plots showing the distribution of RNA transcript quality control metrics, individual cells are shown. **D)** Violin plots showing the proportion of mitochondrial-encoded RNA (left) and ribosomal RNA (right), individual cells are shown.

**Table 2:**
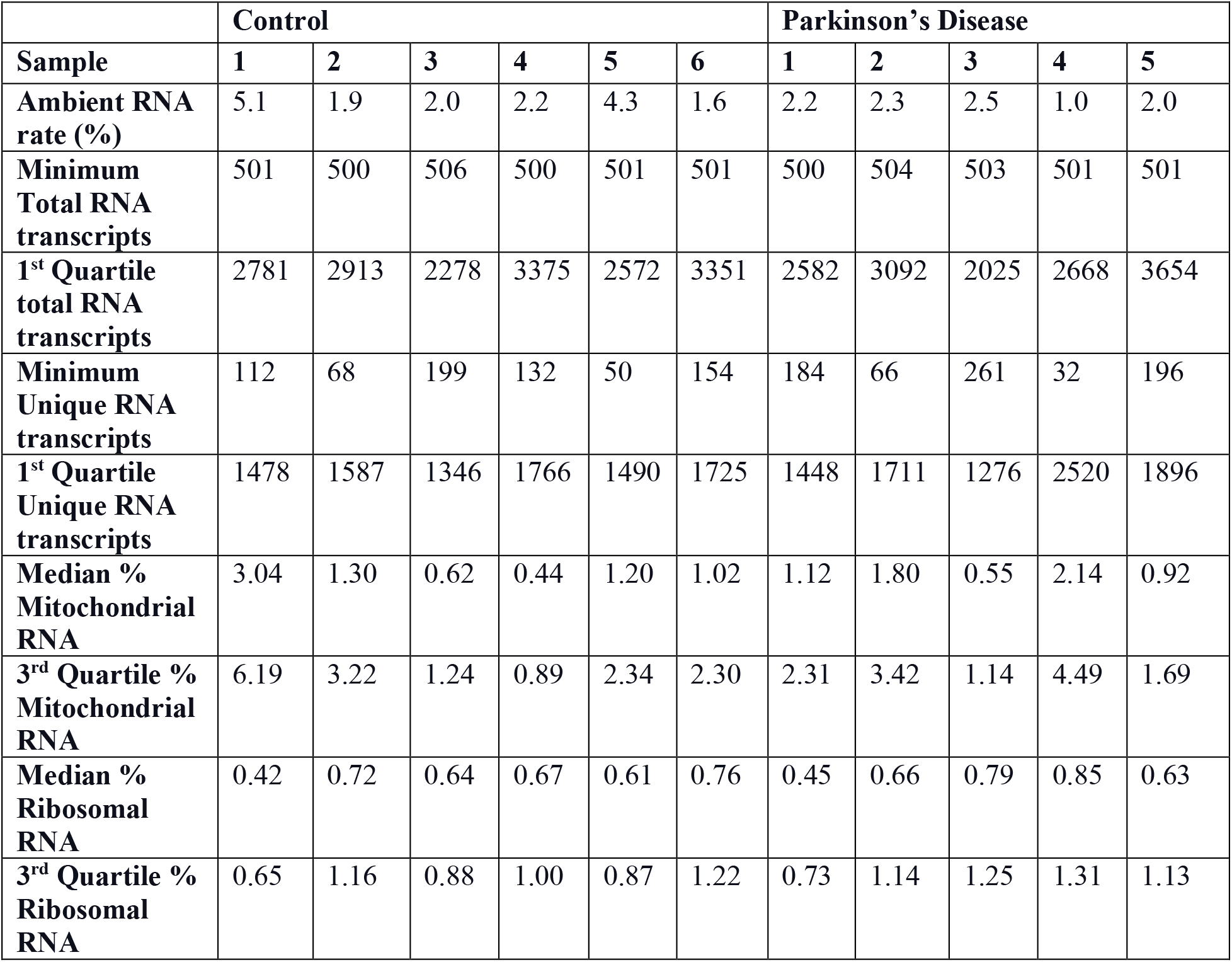
Selected quality control measurements across all samples.

|  | **Control** |  |  |  |  |  | **Parkinson's Disease** |  |  |  |  |
| --- | --- | --- | --- | --- | --- | --- | --- | --- | --- | --- | --- |
| **Sample** | **1** | **2** | **3** | **4** | **5** | **6** | **1** | **2** | **3** | **4** | **5** |
| **Ambient RNA rate (%)** | 5.1 | 1.9 | 2.0 | 2.2 | 4.3 | 1.6 | 2.2 | 2.3 | 2.5 | 1.0 | 2.0 |
| **Minimum Total RNA transcripts** | 501 | 500 | 506 | 500 | 501 | 501 | 500 | 504 | 503 | 501 | 501 |
| **1<sup>st</sup> Quartile total RNA transcripts** | 2781 | 2913 | 2278 | 3375 | 2572 | 3351 | 2582 | 3092 | 2025 | 2668 | 3654 |
| **Minimum Unique RNA transcripts** | 112 | 68 | 199 | 132 | 50 | 154 | 184 | 66 | 261 | 32 | 196 |
| **1<sup>st</sup> Quartile Unique RNA transcripts** | 1478 | 1587 | 1346 | 1766 | 1490 | 1725 | 1448 | 1711 | 1276 | 2520 | 1896 |
| **Median % Mitochondrial RNA** | 3.04 | 1.30 | 0.62 | 0.44 | 1.20 | 1.02 | 1.12 | 1.80 | 0.55 | 2.14 | 0.92 |
| **3<sup>rd</sup> Quartile % Mitochondrial RNA** | 6.19 | 3.22 | 1.24 | 0.89 | 2.34 | 2.30 | 2.31 | 3.42 | 1.14 | 4.49 | 1.69 |
| **Median % Ribosomal RNA** | 0.42 | 0.72 | 0.64 | 0.67 | 0.61 | 0.76 | 0.45 | 0.66 | 0.79 | 0.85 | 0.63 |
| **3<sup>rd</sup> Quartile % Ribosomal RNA** | 0.65 | 1.16 | 0.88 | 1.00 | 0.87 | 1.22 | 0.73 | 1.14 | 1.25 | 1.31 | 1.13 |

To further visualize the data and determine thresholds for filtering, scRNAbox computes the unique RNA transcripts and total counts of RNA for each cell (**Figure 2C**). Cells with too few unique RNA transcripts are only ambient RNA, membrane fragments, or damaged/dying cells, and these barcodes should be removed. The range of unique transcripts varies across species, tissue types, and sample preparations. The distribution of unique RNA transcripts and total RNA varied across the 11 samples; however, the lowest quartile (1^st^ quartile) value was above 1000 in both measures for all samples, indicating that a stringent threshold for good quality cells will retain a large sample size (**Table 2**). Finally, the percentage of mitochondrial and ribosomal RNA transcripts are calculated (**Figure 2D**). A high proportion of mitochondrial-encoded RNA indicates that the mitochondria are damaged within that cell, indicating that the cell is likely dying. In most cases, researchers will remove these cells. Ribosomal RNA genes encode proteins for ribosomal machinery and indicates a high level of translational activity in the cell. Like cell cycle state, elevated levels of ribosomal proteins could later impact clustering results; however, both may also represent biologically relevant signals that researchers may wish to retain and further explore. As expected from nuclear sequencing, the percentage of mitochondrial-encoded genes was low across all samples (**Table 2**).

### ScRNAbox applies quality control filters and integrates samples

In step 3, we applied the filtering criteria used by Samjic et al. (14); we did not adjust for ambient RNA contamination or regress cell cycle genes. We removed unwanted barcodes as described above, applying filters for minimum unique RNA transcripts (>1000), minimum total RNA transcripts (>1500), and maximum percent mitochondria and ribosomal RNA (<10) (**Figure 3A**). Additionally, we removed mitochondrial-encoded and ribosomal genes. After applying these filters, we retained between 2,442 and 6,153 cells per sample (**Table 3**). In Step 4, we leveraged DoubletFinder to predict doublets using default parameters and 25 PCs and defined the expected doublet rate for each sample based on the number of recovered cells from the CellRanger pipeline (**Figures 3B and 3C**; **Table 3**). The DoubletFinder algorithm requires that UMAP dimensional reduction is performed prior to analysis. We performed dimensional reduction using 25 PCs and 65 nearest neighbours. After removing predicted doublets, 44,538 cells remained across all samples. In total, 9,460 cells (17.52%) were filtered from the dataset (**Table 3**).

**Figure 3:**
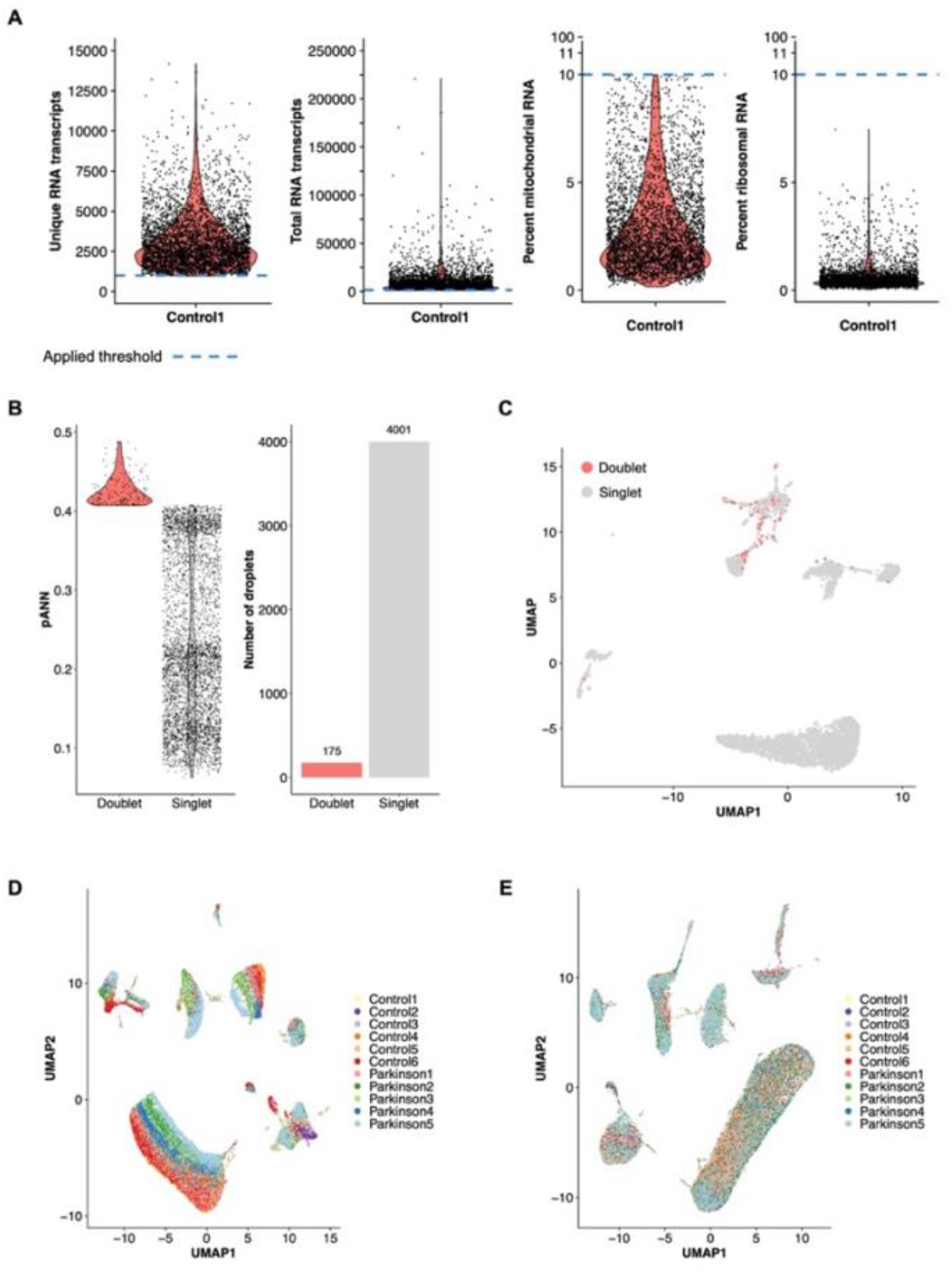
scRNAbox produces visualizations of filter application, doublet detection and data integration. **A)** Violin plots visualizing the distribution of quality control metrics after filtering according to user-defined thresholds, for snRNAseq midbrain sample *Control 1*. **B)** Results of doublet detection with DoubletFinder (21). Left: violin plot displaying the distribution of the proportion of artificial nearest neighbours (pANN) across singlets and doublets for *Control 1*. Right: a bar plot of the number of predicted singlets and doublets for *Control 1*. **C)** Uniform Manifold Approximation Projection (UMAP) plots coloured by droplet assignments (singlet or doublet) for *Control 1*. **D)** UMAP of merged snRNAseq midbrain samples (six Control and five PD) coloured by sample identity. **E)** UMAP of the same data after integration using Seurat label transfer, coloured by sample identity.

**Table 3:** Unique barcode counts at different stages of data processing.

| Sample | Number of Unique Barcodes | Cell count after filtering | Expected doublet rate (%) | Number of doublets detected | Cell count after doublet removal |
| --- | --- | --- | --- | --- | --- |
| Control 1 | 5513 | 4176 | 4.2 | 175 | 4001 |
| Control 2 | 5508 | 4553 | 4.2 | 191 | 4362 |
| Control 3 | 2856 | 2490 | 2.3 | 57 | 2433 |
| Control 4 | 5347 | 4952 | 4.0 | 198 | 4754 |
| Control 5 | 3564 | 3196 | 2.7 | 86 | 3110 |
| Control 6 | 7174 | 6153 | 5.3 | 326 | 5827 |
| PD 1 | 2899 | 2442 | 2.3 | 56 | 2386 |
| PD 2 | 7169 | 6388 | 5.3 | 339 | 6049 |
| PD 3 | 4560 | 3784 | 3.4 | 129 | 3655 |
| PD 4 | 2820 | 2307 | 2.3 | 53 | 2254 |
| PD 5 | 6588 | 6007 | 5.0 | 300 | 5707 |
| Total | 53998 | 46448 | NA | 1910 | 44538 |

**Abbreviations:** PD, Parkinson's disease.

Finally, after processing each individual sample, we combined all samples into one data object to facilitate integrated analysis. In Step 5, users have the option to either merge (**Figure 3D**) or integrate (**Figure 3E**) the data. We proceeded with downstream analyses of the midbrain dataset using the integrated data object, which facilitates the identification of cell types that are consistent across samples (22).

### ScRNAbox provides tools to optimize clustering and facilitate annotation

In Step 6, we performed clustering on the integrated dataset to eventually identify distinct cell types. We clustered the cells using the 4000 most variably expressed features and 25 PCs, maintaining the parameters used by Smajic et al. (14). We used 30 neighbours to construct the shared nearest neighbour graph input into the Louvain network detection algorithm and performed clustering on a range of clustering resolutions (**Supplementary Figure S2A**). To evaluate the reproducibility of clusters identified at each resolution, we calculated the ARI between clustering pairs at each resolution across 25 replications (23). The ARI at a clustering resolution of 0.05 and 0.2 were both 1.00 and the ClusTree plot suggested high stability (**Supplementary Figures S2B and S2C**). Thus, we used a clustering resolution of 0.2, which identified 14 clusters, to annotate the major cell types (**Figure 4A**).

**Figure 4.**
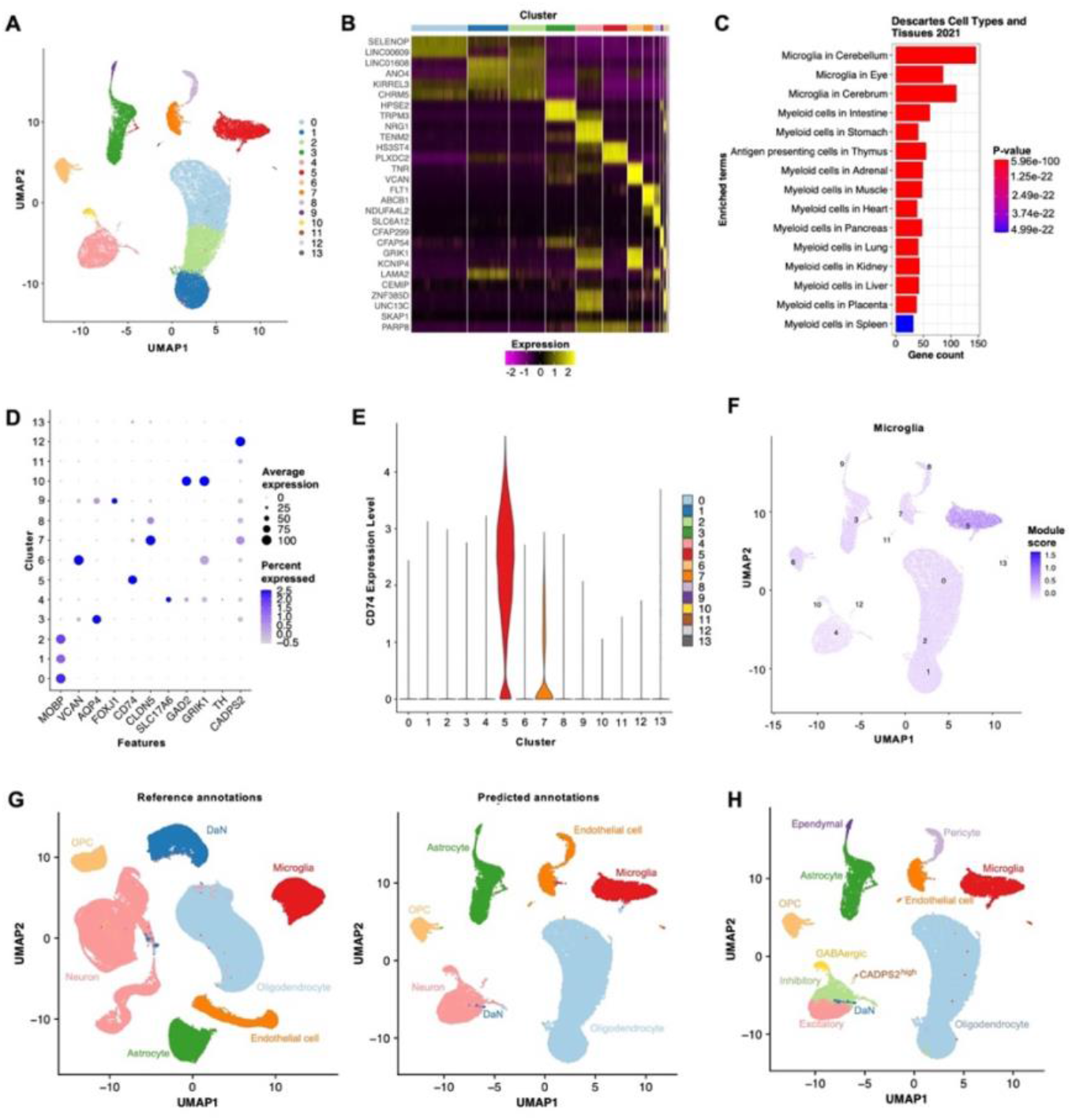
scRNAbox performs clustering to identify cells type groups and provides tools for cluster annotation. **A)** Uniform Manifold Approximation Projection (UMAP) plots showing clusters identified by Louvain network detection with a resolution of 0.2, coloured by cluster index. The UMAP was generated from the 11 integrated snRNAseq midbrain samples. **B)** Heatmap of the top 3 upregulated marker genes for each cluster in A. **C)** Bar chart showing the top 15 cell types in the *Descartes Cell Types and Tissue* library identified by GSEA of the marker genes for cluster 5. **D)** Dot plot showing expression of cell type markers defined by Smajic et al. for each cluster at resolution 0.2. The cell type markers are as follows: oligodendrocytes: *MOBP*; oligodendrocyte precursor cells (OPC): *VCAN*; astrocytes: *AQP4*; ependymal cells: *FOXJ1*; microglia: *CD74*; endothelial cells: *CLDN5*; pericytes: *GFRB*; excitatory neurons: *SLC1746*; inhibitory neurons: *GAD2*; GABAergic neurons: *GAD2* and *GRIK1*; dopaminergic neurons (DaN): *TH*. PD specific DaN subgroup; *CADPS2*. **E)** Violin plot showing expression levels in each cluster across individual cells for the microglia marker *CD74*. **F)** UMAP showing the module score for the microglia gene marker list. The module score is an aggregated expression of known marker genes (20). **G)** Left: UMAP of clustered and annotated reference Seurat object; snRNAseq of midbrain tissue produced by Kamath et al. (31), coloured by cell type. Using the Seurat label transfer approach, the reference data was used to predict cell types in the query data (11 snRNAseq midbrain samples from Smajic et al. (14)). Right: UMAP of the label transfer predictions for each cell, coloured by predicted cell type. **H)** UMAP of the 11 integrated samples with the applied final cell type annotation, coloured by cell type.

In Step 7, we applied the three cluster annotation tools within the scRNAbox pipeline to identify the cell types. Using Tool 1, we identified the top markers for each cluster (**Figure 4B**) and subjected these to genes GSEA using the EnrichR R package. As an example, the *Descartes Cell Types and Tissues 2021* library GSEA suggested that cluster 5 are microglia (**Figure 4C**). For Tool 2, we profiled the expression of known marker genes, using the marker genes identified by Samjic et al. to annotate their clusters: oligodendrocytes: *MOBP*; oligodendrocyte precursor cells (OPC): *VCAN*; astrocytes: *AQP4*; ependymal cells: *FOXJ1*; microglia: *CD74*; endothelial cells: *CLDN5*; pericytes: *GFRB*; excitatory neurons: *SLC1746*; inhibitory neurons: *GAD2*; GABAergic neurons: *GAD2* and *GRIK1*; dopaminergic neurons (DaN): *TH*. Except for clusters 11 and 13, we found that each cluster showed elevated expression for at least one marker gene (**Figure 4D**). ScRNAbox also allows expression profiling of known marker genes through a violin plot. For instance, we explored the expression of *CD74* across clusters and found that cluster 5 showed elevated expression of this gene, further suggesting that this cluster consists of microglia (**Figure 4E**). Next, we computed the module scores for custom gene marker lists (**Supplementary Table S1**). The module score for the microglia gene set was highest in cluster 5 (**Figure 4F**). Using Tool 3, we predicted cell types using a labelled Seurat object generated from snRNAseq midbrain data published by Kamath et al. (31) (**Figure 4G**).

Performing cluster annotations at a clustering resolution of 0.2 allowed us to identify the major cell types expected in the human midbrain. However, to further classify the neurons into subtypes, we repeated Step 7 at a clustering resolution of 1.5, as used by Smajic and colleagues (14). We subjected the 33 clusters identified to marker GSEA and profiled the expression of known marker genes and cell type marker gene lists (**Supplementary Figures S3-6**). We identified each of the expected neuronal subtypes, including a cluster of CADPS2^high^ DaNs identified by Smajic et al., resulting in 12 cell types for our final annotation (**Figure 4H; Supplementary Figure S6D**).

### ScRNAseq efficiently calculates differential gene expression and facilitates pathway analysis

In Step 8, scRNAbox calculates DGE by two different methods, cell-based using MAST (26) and sample-based using DESeq2 (28). We first added metadata to the Seurat object and classified each sample as either “Control” or “PD”, allowing us to define our desired contrasts. Next, we computed DGE between PD and controls for all cells together and for each cell type individually. Cell-based DGE results in fewer DEGs with L2FC greater than 1.0 but more genes significant after p-value adjustment for multiple comparisons (**Figure 5A and 5B; Supplementary Figure S8 and S9; Supplementary Table S2 and S3**). For example, cell-based-DGE identified 13 DEGs (p-value < 0.05; L2FC > 1) between PD and controls for microglia, while pseudo-bulk with DESeq2 identified 1,030 DEGs at the same significance threshold and for the same cell type (**Figure 5A and 5B**). Indeed, the sample-based-DGE identified a higher number of DEGs across all cell types compared to MAST, except for DaNs (sample-based = 82 DEGs; cell-based = 111 DEGs) (**Figure 5C and 5D**). Another benefit of using multiple statistical frameworks for computing DGE is the ability to identify consensus signals. Particularly, the DEGs that maintain significance after correction for multiple hypothesis testing by multiple statistical frameworks may be of highest interest to investigators (**Figure 5E**). Finally, the DGE data tables produced by the scRNAbox pipeline can be used to perform gene enrichment pathway analysis and explore the contribution of different cell types to perturbed pathways. As an example, we performed GO analysis for biological processes using significant DEGs (p-value < 0.05 and L2FC > 1) identified by sample-based DGE (1,366 genes) and cell-based DGE (7 genes), comparing all cells between PD and control subjects using the *ClusterProfiler* R package (29). We then selected the top 5 most significantly enriched pathways for each method and looked at the gene contribution and pathway significance for each GO term across cell types (**Figure 5F and 5G**). Interestingly, both DGE methods suggested perturbed pathways related developmental and neuro-anatomical changes in the PD midbrain.

**Figure 5.**
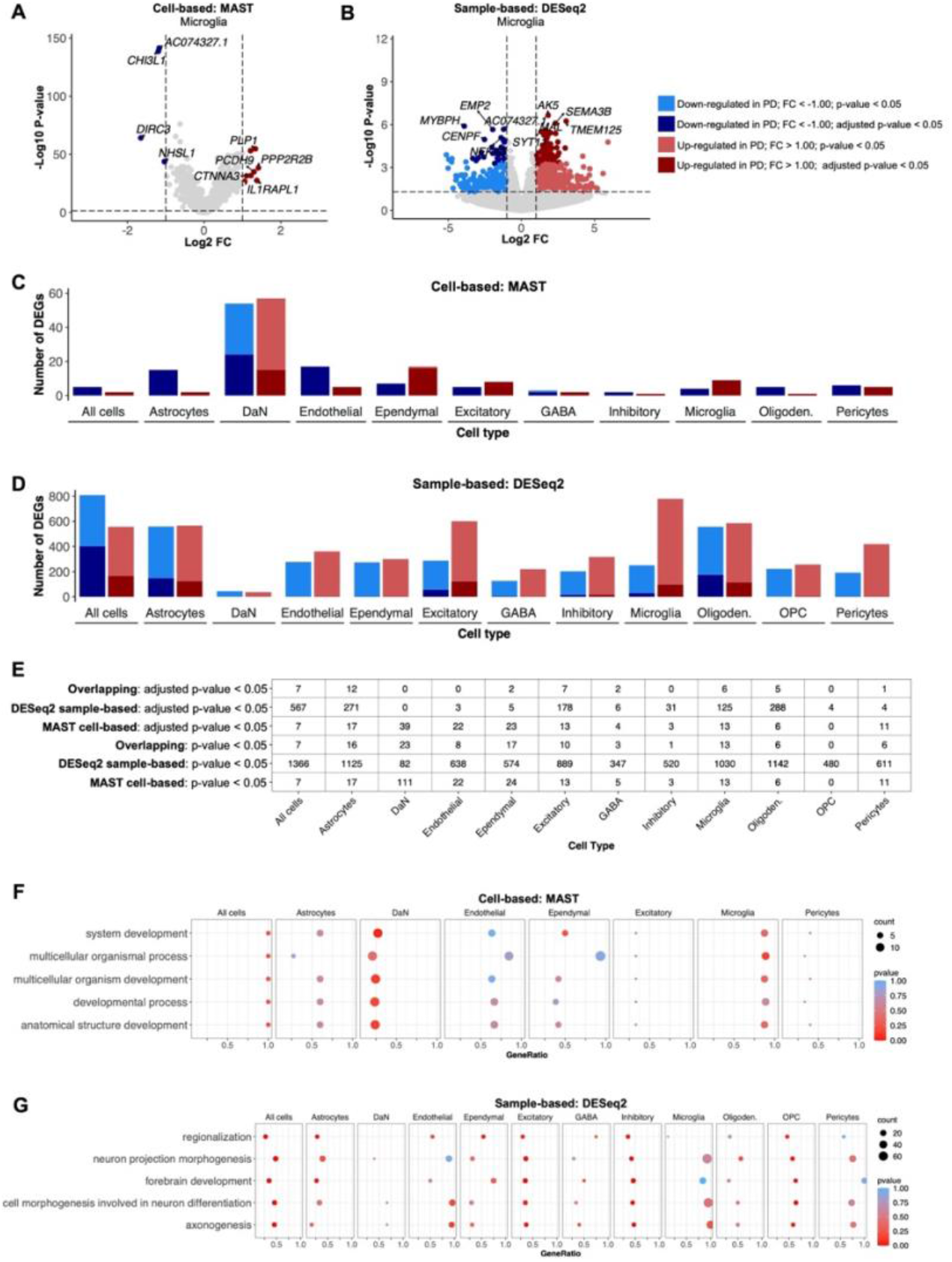
scRNA calculates differential gene expression (DGE) using multiple statistical frameworks. ScRNAbox computes DGE using two distinct data preparations: 1) using cells as replicated and the MAST statistical framework (26) and 2) using samples as replicates (pseudo-bulk) and the DESeq2 statistical framework (28). **A)** Volcano plot showing cell-based DGE results identified by MAST, between Parkinson’s disease (PD) and control subjects for microglia. **B)** Volcano plot showing sample-based DGE identified by DESeq2 between PD and control subjects for microglia. **C, D)** Bar chart showing the number of DEGs identified with a log 2 fold-change (L2FC) (−1 < and > 1) and p-values < 0.05. Bonferroni adjusted p-values < 0.05 are indicated by the darker shade. **C)** Cell-based DGE using MAST. **D)** Sample-based DGE using DESeq2 **E)** Number of differentially expressed genes (DEG) identified by cell-based DGE-MAST, sample-based DGE-DESeq2, or both frameworks across all cell types. Only DEGs with L2FC (−1 < and > 1) are included. **F, G)** Bar chart showing the top 5 GO terms for GO-Biological Processes calculated for all cell types together. DEGs with p-values < 0.05 and L2FC (−1 < and > 1) were used as the input for gene set enrichment analysis (GSEA). The gene ratio, gene count, and p-value of the five terms in each cell type are shown. **F)** GO analysis of DEGs identified by cell-based DGE across all cell types. The missing cell types did not have enough DEGs for GSEA analysis to return results and were not plotted. **G)** GO analysis of DEGs identified by sample-based DGE across all cell types.

### ScRNAbox effectively demultiplexes cells with Hashtag feature labels

In addition to standard scRNAseq data, scRNAbox can be used to analyze multiplexed samples, whereby each subject is tagged with a unique HTO, pooled, and then captured and sequenced together. Cell hashtagging can reduce the cost of scRNAseq by a factor of the number of samples multiplexed; however, additional steps are required to bioinformatically assign each cell back to its sample of origin. In Step 4, scRNAbox provides the option for users to demultiplex cells based on the expression of sample specific HTOs. To demonstrate, we analyzed a scRNAseq dataset of PBMCs from 8 subjects collected by Stoeckius et al. (16). In Step 0, we selected to run the “HTO” analysis track and proceeded to run Steps 1-3 using the same analytical parameters that Stoeckius et al. used to process their data. At Step 4, instead of running doublet detection with DoubletFinder, scRNAbox uses the Seurat *MULTIseqDemux* function to assign cells back to their sample-of-origin based on HTO expression (17). ScRNAbox produces multiple figures to visualize the enrichment of HTOs across samples. Upon examining the expression levels of each HTO label across samples, we observed that cells with a distinct expression for a given HTO are assigned to the matching sample (**Figures 6A-C**). Barcodes with multiple HTO labels are detected as doublets, as these likely represent two cells that were sequenced together. Negative cells have too low of a level of any HTO tag to be accurately assigned. The doublet group has about twice as many RNA transcripts per cell compared to the cells that were assigned to an individual sample, suggesting that the predicted doublets are likely true doublets (**Figure 6D**). We conclude that scRNAbox pipeline can accurately demultiplex samples with HTO tags.

**Figure 6.**
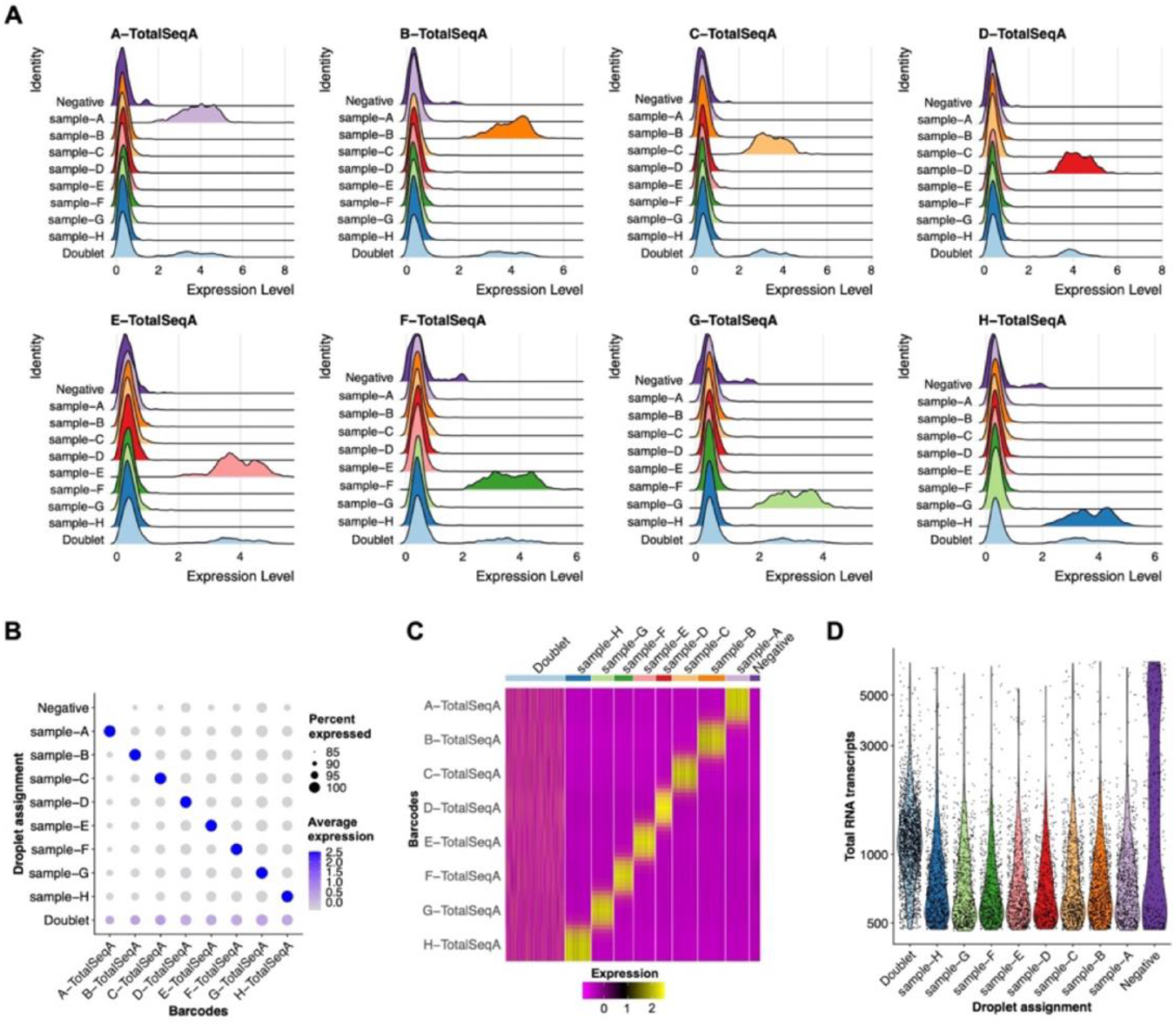
scRNAseq effectively demultiplexes HTO samples and detects doublets. The expression matrices of sample-specific oligonucleotide conjugated antibodies (HTO) are used to demultiplex samples and identify doublets (17). The enrichment of barcode labels across sample assignments are visualized at the cellular and sample level. **A)** Ridge plots (stacked density plots) showing the expression each HTO tag (X-total-seq) expression in each assigned sample. **B)** Dot plot showing the expression level (colour intensity) and proportion of cells (dot size) expressing each HTO in each assigned sample. **C)** Heatmap showing expression levels of each HTO tag in each assigned sample. **D)** Violin plot showing the distribution of total RNA transcripts across sample assignments.

## Conclusions

Here, we introduce ScRNAbox, a comprehensive end-to-end pipeline designed to streamline the processing and analysis of single-cell transcriptomic data. ScRNAbox responds to the pressing demand for a user-friendly, HPC solution, bridging the gap between the growing computational demands of scRNAseq analysis and the coding expertise required to meet them. ScRNAbox empowers researchers, regardless of coding experience, to unlock the full potential of HPC clusters. By automating and optimizing the entire scRNAseq analysis workflow, it facilitates the processing of numerous samples while seamlessly scaling to meet user needs. The stepwise execution of ScRNAbox provides researchers with fine-grained control over parameters and manual cell annotations, ensuring reproducibility and customizability at every stage. The pipeline contains a rich array of functionalities, enabling cell type annotation, differential gene expression analysis, and efficient cell demultiplexing using Hashtag feature labels.

While ScRNAbox offers an efficient solution for scRNAseq data analysis, it does come with certain limitations. Primarily tailored for sequencing alignment from 10X data and focused on differential gene expression analysis, ScRNAbox does not encompass trajectory analysis, cell-to-cell networks, or other downstream analytical methods. Nonetheless, it equips users with final and intermediate data objects that seamlessly integrate into external packages for advanced analyses.

Using ScRNAbox, we conducted a comprehensive analysis of snRNAseq data from PD and control midbrains, provided by Smajic et al. We elucidate each analysis step and demonstrate the remarkable alignment of our annotations with the original cell types and proportions. Additionally, we perform a comparative analysis of two DGE methods, shedding light on overlapping genes between the two approaches.

Our open-source, modular code provides a versatile foundation for users to customize and expand. We encourage researchers to harness the flexibility of ScRNAbox, introducing alterations, additional options, or their preferred downstream analyses. With ScRNAbox, we aspire to simplify the intricacies of scRNAseq analysis, inviting an extended community of researchers to embark on novel and thoughtful explorations of single-cell transcriptomics.

## Supporting information

BioRender publication license

## Acknowledgements

This work was supported by the Canadian Institute of Health Research [FDN – 154301 to EAF]; and the Michael J. Fox Foundation [MJFF-020696 to EAF]. Additionally, RAT received funding through the McGill Healthy Brains for Healthy Lives (HBHL) Postdoctoral Fellowship and Molson NeuroEngineering Fellowship. MRF is supported by a CIHR Canada Graduate Scholarships-Master’s Award, a Fonds de recherche Santé Québec Master’s Award (BF1 – 334616), and a Brain Canada Rising Stars Award. EAF is supported by a Fonds d’Accéleration des Collaborations en Santé (FACS) grant from CQDM/MEI, a Canada Research Chair (Tier 1) in Parkinson’s disease. SMKF received funding from Brain Canada and The Montreal Neurological Institute-Hospital. The authors acknowledge Indra Roy for preparation of the annotated reference dataset. We thank Malosree Maitra and Moein Yaqubi for discussions about scRNAseq analysis.

## Contributions

Conception RAT, MRF, SA, and SMKF conceived the project. RAT designed the pipeline. RAT, MRF, and SA wrote R scripts. SA created the HPC pipeline and wrote bash submission scripts. MRF and SA implemented the R scripts into the HPC pipeline. SA created the GitHub site. MRF wrote the documentation. RAT, MRF, and SA tested the pipeline. MRF and SA debugged the code. MRF conducted the analysis. MRF produced the figures and tables. RAT and MRF wrote the manuscript with input and final approval from all authors. EAF provided funding. SMKF supervised the project.

