## Supplementary Material for "ScRNAbox: Empowering Single-Cell RNA Sequencing on High Performance Computing Systems"

**Supplemental Materials**

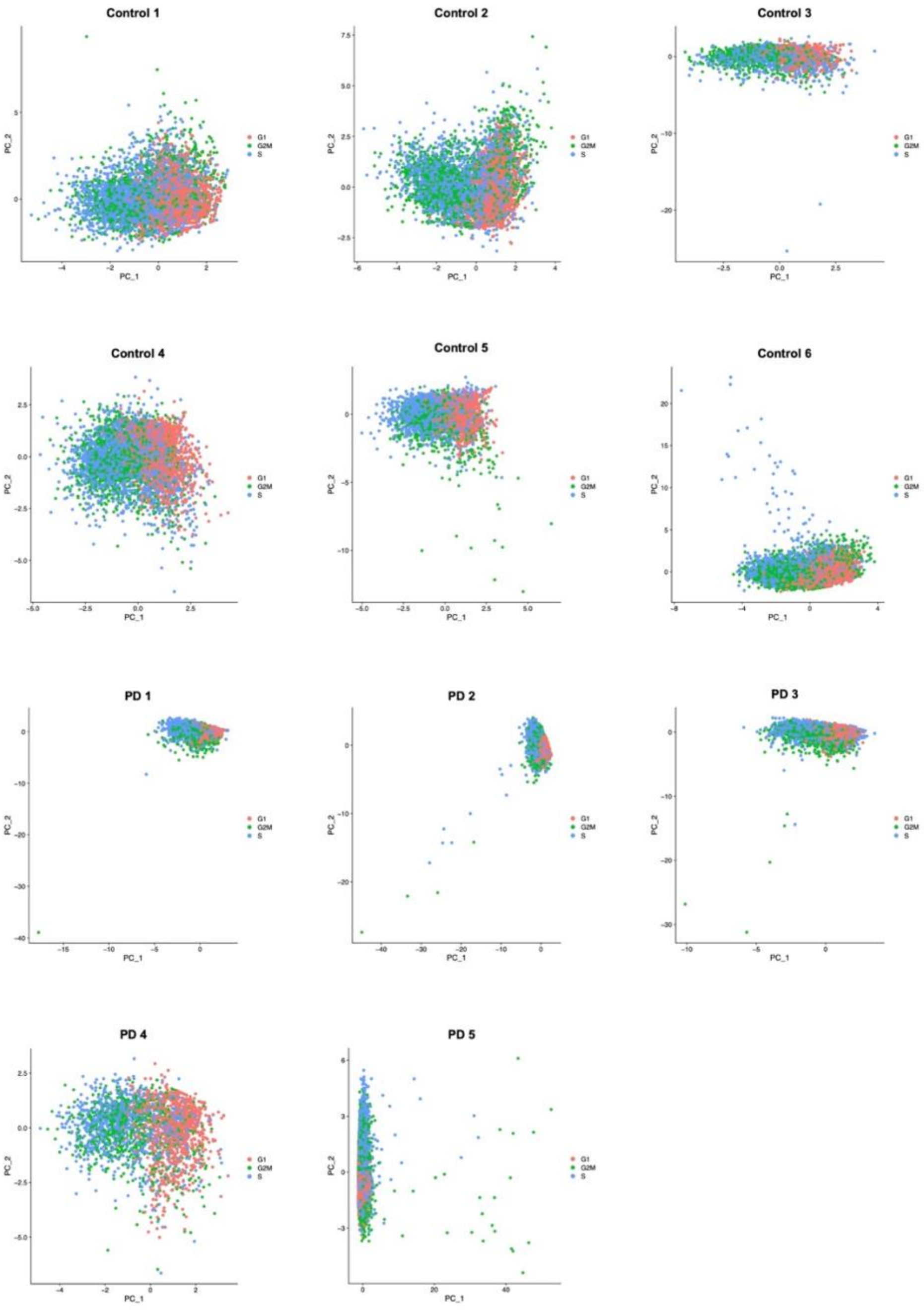


**Figure S1. Cell cycle phases analysis.** Scatter plots of Principal Component Analysis of cells from each of the 11 samples from the midbrain dataset. Cells are coloured by cell cycle phase calculated using the Seurat S and G2M reference genes and show the impact of cell cycle genes on distribution of cells in PCA space. The sample names are shown in the plot titles. PCA plots are produced in Step 2 of the scRNAbox pipeline.

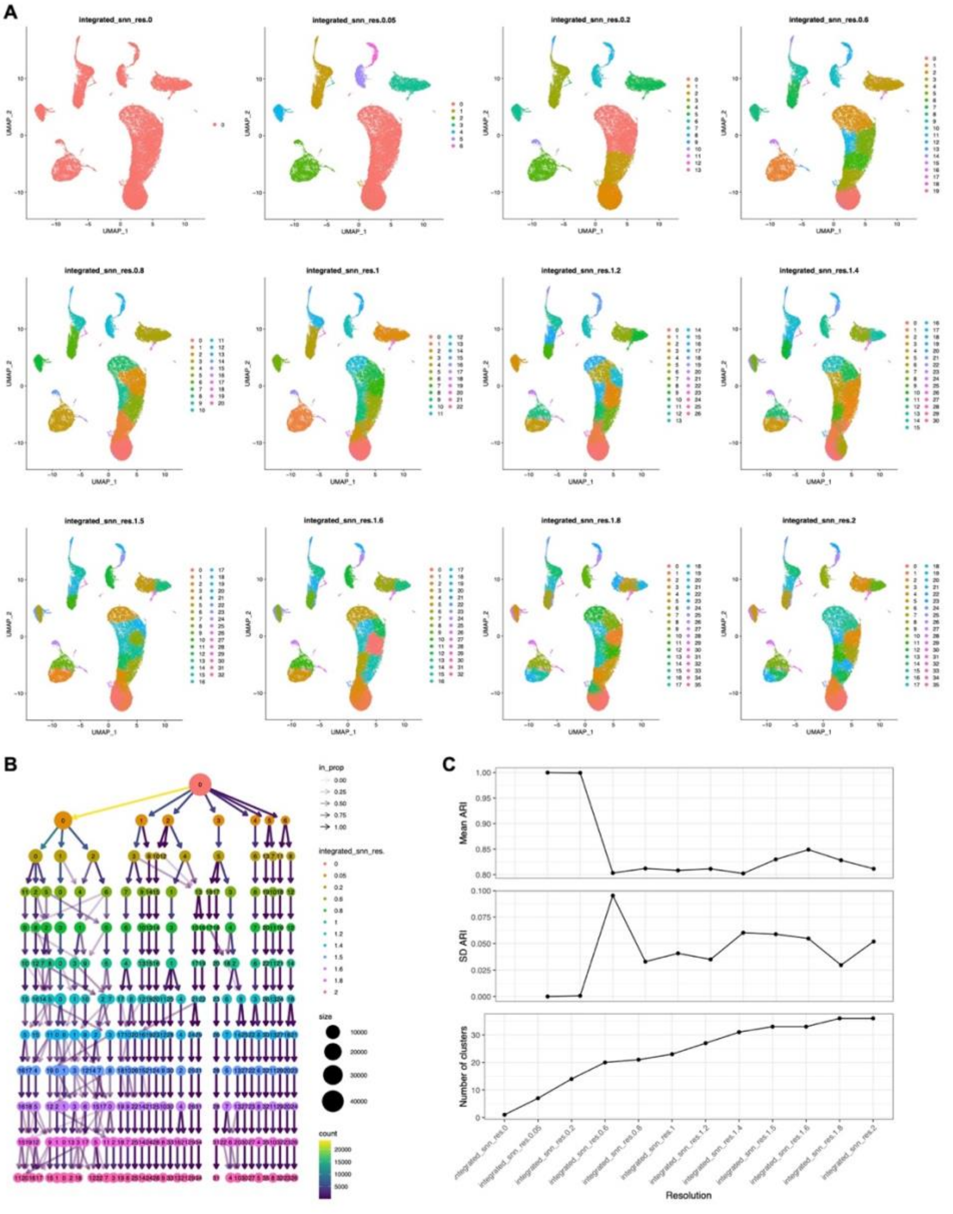


**Figure S2. Clustering across a range of resolutions.** In Step 6, scRNAbox allows users to define a list of clustering resolutions in the execution parameters to investigate their data at different granularities. For the midbrain dataset, we performed clustering at 12 resolutions: 0, 0.05, 0.2, 0.6, 0.8, 1.0, 1.2, 1.4, 1.5, 1.6, 1.8, and 2.0. **A)** Uniform manifold approximation and projection (UMAP) plots for each resolution, coloured by cluster index. Resolutions are indicated in the plot titles. **B)** A ClustTree plot visualizing inter-cluster dynamics at varying cluster resolutions. **C)** To evaluate the reproducibility of each cluster, scRNAbox performs Louvain clustering multiple times at each clustering resolution, while shuffling the order of the nodes in the graph for each iteration. The mean (top panel) and standard deviation (SD; middle panel) of the adjusted rand index (ARI) between clustering pairs at each clustering resolution was then calculated (25 iterations).

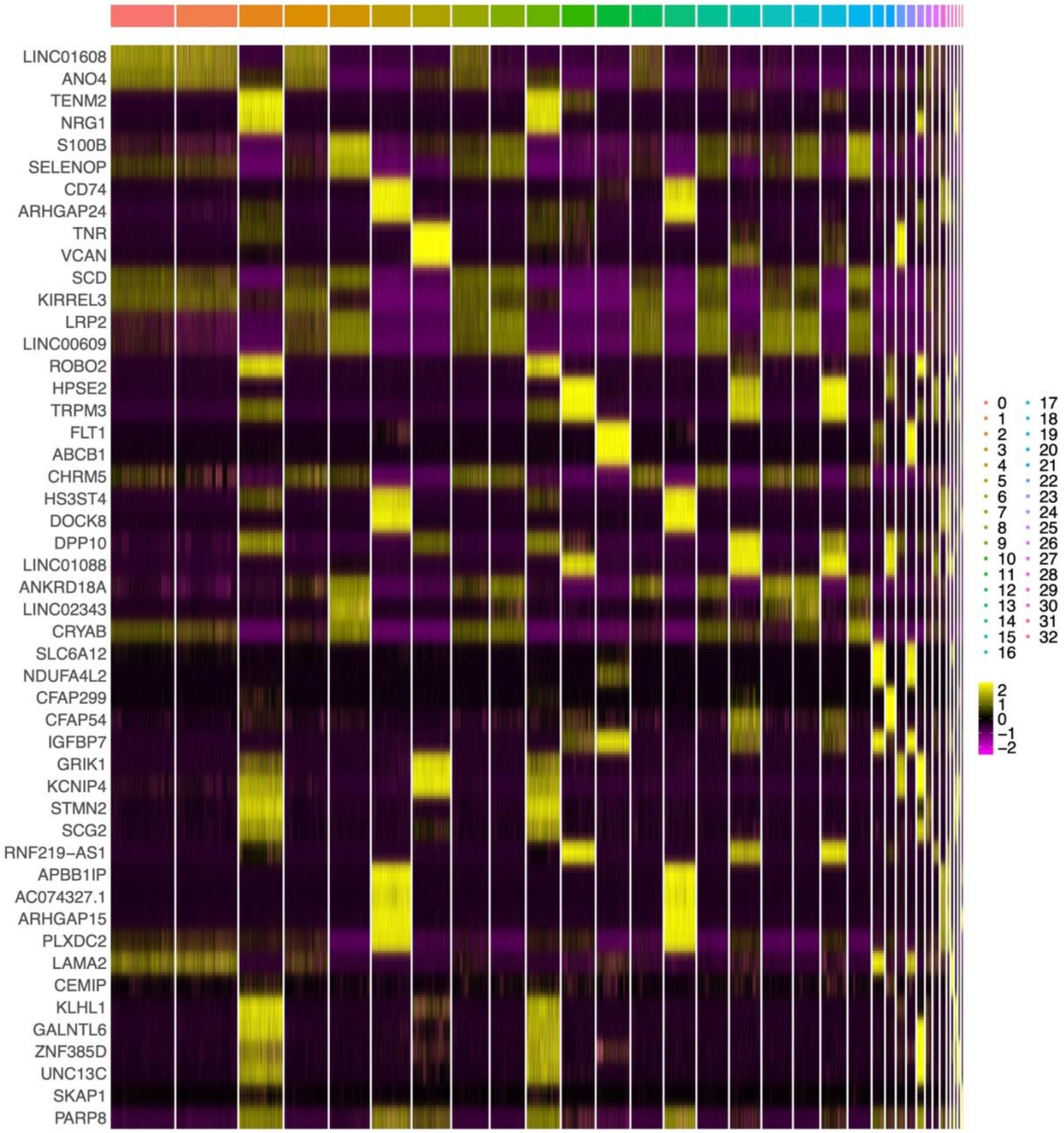


**Figure S3. Visualizing the expression of the top marker genes for each cluster.** Heat map of the top cluster markers in the midbrain dataset at a clustering resolution of 1.5, which identified 33 clusters.

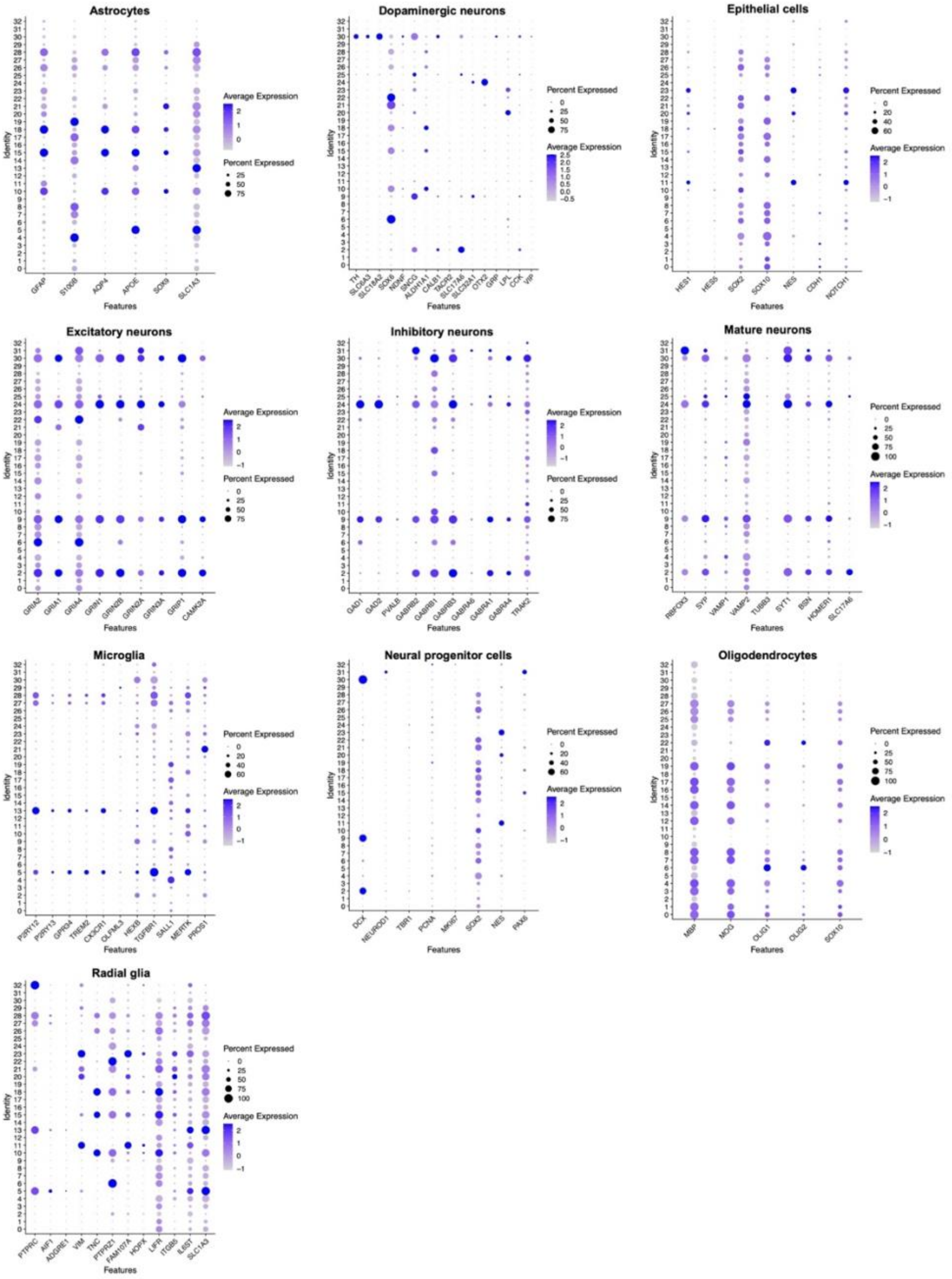


**Figure S4. Profiling the expression of known marker genes at the cluster level.** Using Step 7 cluster annotation tool 2, dot plots of custom gene list were produced to identify cell types in the midbrain dataset at a clustering resolution of 1.5, which identified 33 clusters. The cell types defined by the marker gene list are indicated in the dot plot titles. Expression levels of each gene (x axis) are shown by intensity and the proportion of cells expressing each gene is shown by dot size for each cluster (y axis).

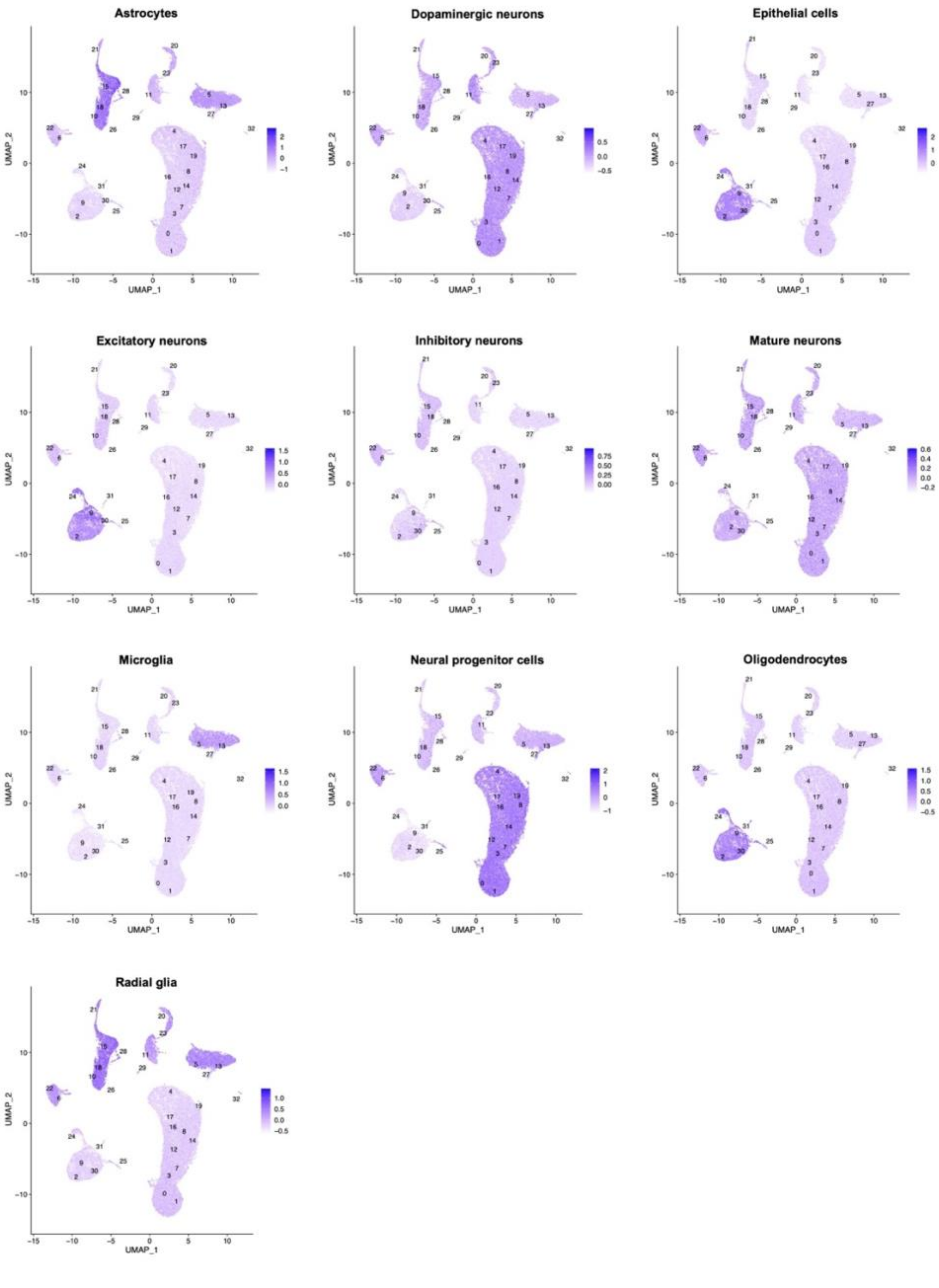


**Figure S5. Visualization of the module scores of known marker genes.** UMAPs of in the midbrain dataset, clustered at resolution of 1.5, the 33 clusters identified are labelled on the plots. Step 7, tool 2 was used to compute the module score (aggregated expression of known marker genes) of common cell types in the human midbrain, cell types are indicated above each plot. The colour intensity corresponds to the module score for each cell.

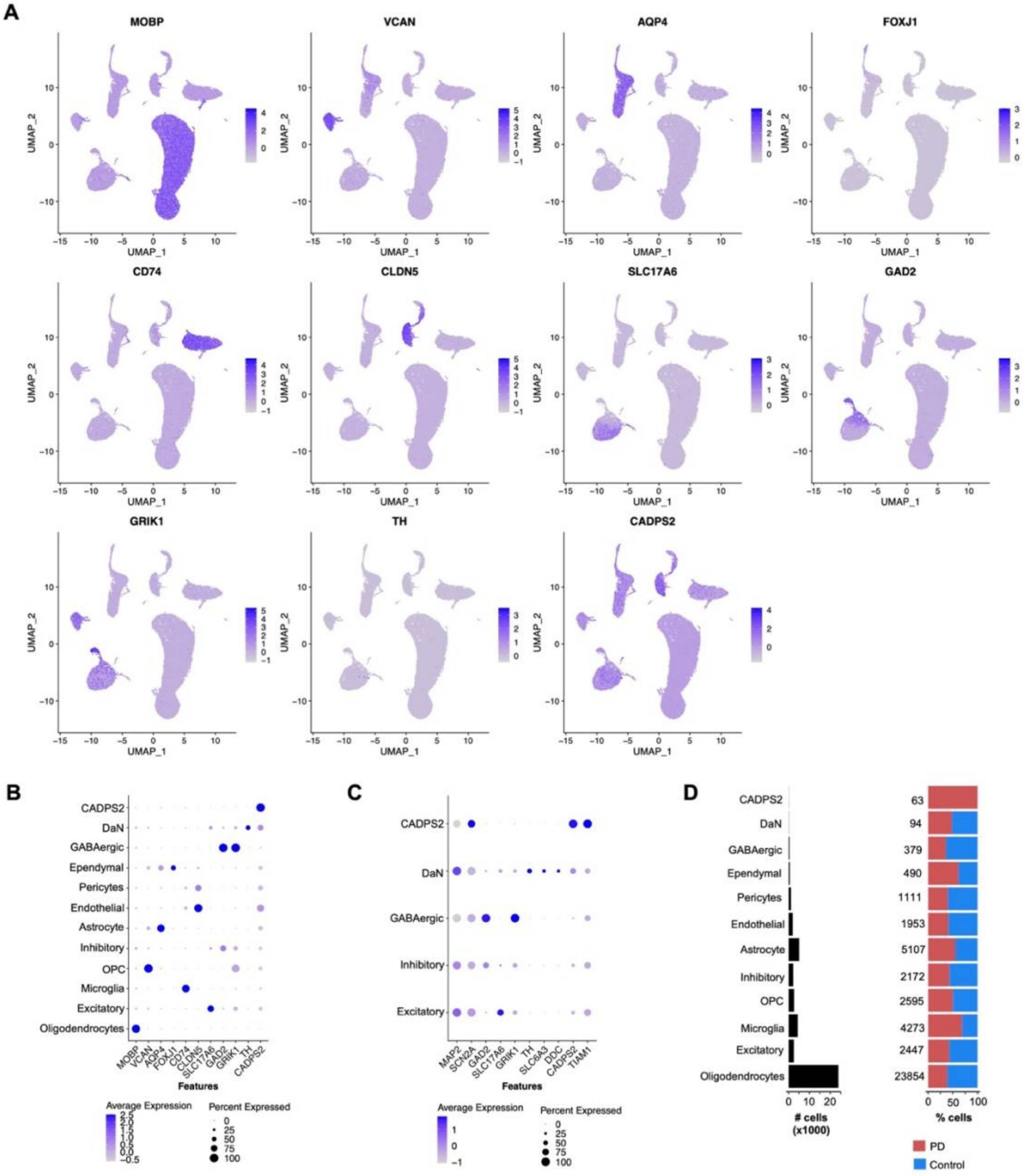


**Figure S6. Profiling the expression of marker genes used by Smajic et al. to define cell types. A)** UMAPs showing the expression level of each cell type marker, the corresponding gene is indicated in the plot titles. **B and C)** Dot plots showing expression levels of marker genes (x axis) in annotated cell type groups (y axis). Dot size indicates the proportion of cells expressing a gene and intensity indicates expression level. **B)** All cells. **C)** Neuronal cells. **D)** Left: the number of cells annotated to each cluster. Right: the proportion of cells belonging to Parkinson's disease (PD) or control subjects.

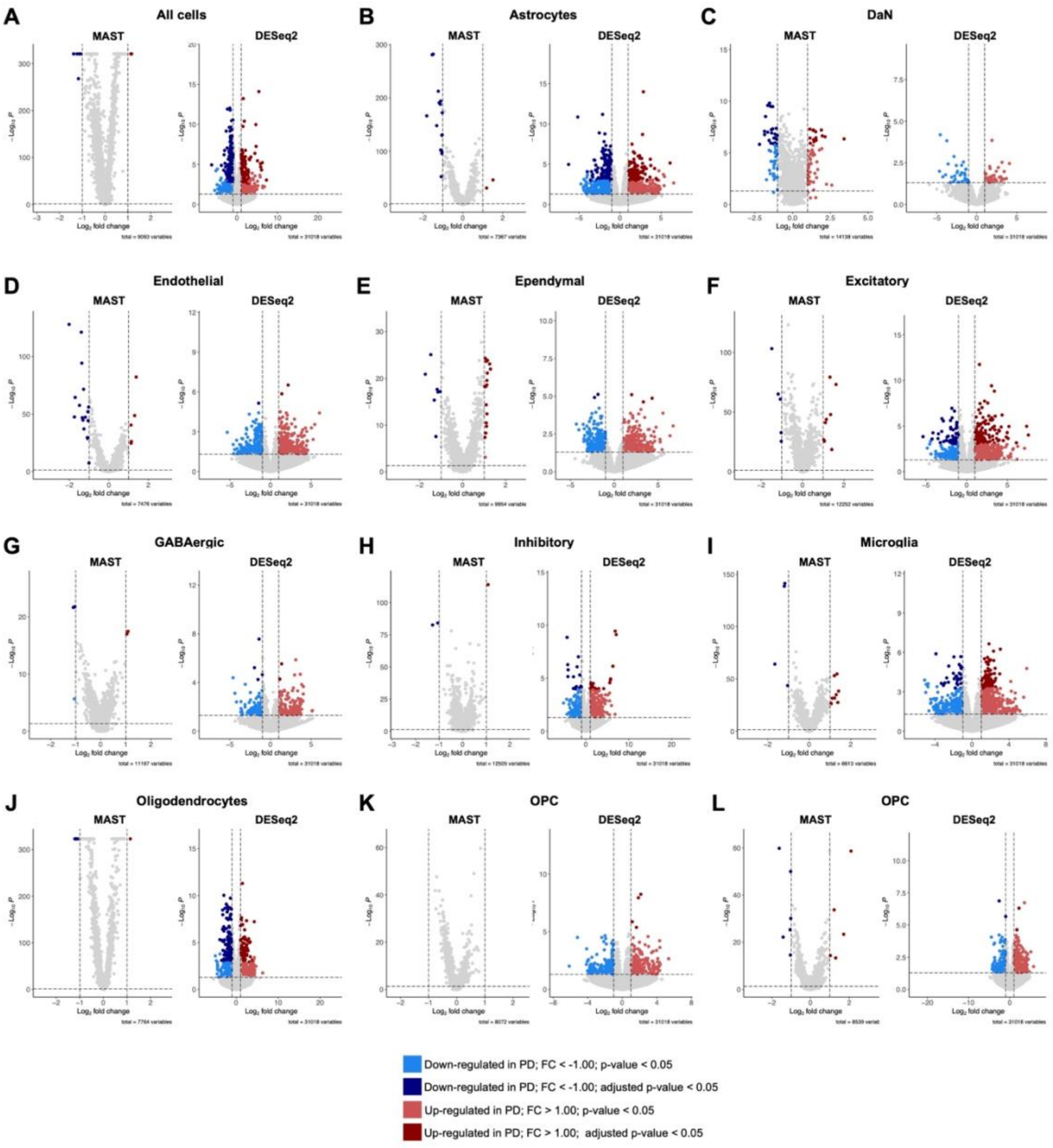


**Figure S7. Differential gene expression (DGE) between Parkinson's disease (PD) subjects and controls across cell types.** DGE between PD subjects and controls across the midbrain dataset using two methods: cell-based, using cells as replicates and the MAST package; and sample-based, using samples as replicates with DESeq2. Volcano plots showing differentially expressed genes were produced for DGE methods the following contrasts: **A)** all cells, **B)** astrocytes, **C)** dopaminergic neurons (DaN), **D)** endothelial cells, **E)** Ependymal cells, **F)** excitatory neurons, **G)** GABAergic neurons, **H)** Inhibitory neurons, **I)** microglia, **J)** oligodendrocytes, **K)** oligodendrocyte precursor cells (OPC), and **H)** Pericytes.

**Table S1. Marker genes for common cell types found in the human midbrain.**

| DaN | NPC | Mature neuron | Excitatory neuron | Inhibitory neuron | astrocyte | Oligoden-drocytes | Radial glia | Epithelial cells | Miroglia |
|---|---|---|---|---|---|---|---|---|---|
| TH | DCX | RBFOX3 | GRIA2 | GAD1 | GFAP | MBP | PTRPC | HES1 | IBA1 |
| SLC6A3 | NEUROD1 | SYP | GRIA1 | GAD2 | S100B | MOG | AIF1 | HES5 | P2RY12 |
| SLC18A2 | TBR1 | VAMP1 | GRIA4 | GAT1 | AQP4 | OLIG1 | ADGRE1 | SOX2 | P2RY13 |
| SOX6 | PCNA | VAMP2 | GRIN1 | PVALB | APOE | OLIG2 | VIM | SOX10 | TREM119 |
| NDNF | MKI67 | TUBB3 | GRIN2B | GABR2 | SOX9 | SOX10 | TNC | NES | GPR34 |
| SNCG | SOX2 | SYT1 | GRIN2A | GABR1 | SLC1A3 | | PTPRZ1 | CDH1 | SIGLECH |
| ALDH1A1 | NES | BSN | GRIN3A | GBRR1 | | | FAM1074 | NOTCH1 | TREM2 |
| CALB1 | PAX6 | HOMER1 | GRIN3 | GABRB2 | | | HOPX | | CX3CR1 |
| TACR2 | | SLC17A6 | GRIP1 | GABRB1 | | | LIFR | | FCRLS |
| SLC17A6 | | | CAMK2A | GABRB3 | | | ITGB5 | | OLFML3 |
| SLC32A1 | | | | GABRA6 | | | IL6ST | | HEXB |
| OTX2 | | | | GABRA1 | | | SLC1A3 | | TGFBR1 |
| GRP | | | | GABRA4 | | | | | SALL1 |
| LPL | | | | TRAK2 | | | | | MERTK |
| CCK | | | | | | | | | PROS1 |
| VIP | | | | | | | | | |

Abbreviations: DaN, dopaminergic neurons; NPC, neural progenitor cells.

**Table S2. Differentially expressed genes (DEG) across cell types identified in the midbrain dataset using the MAST statistical framework.** DEGs with p-value < 0.05 and log2 fold-change > 1.00 are shown.
*See Supplemental_Table2.xlsx*

**Table S3. Differentially expressed genes (DEG) across cell types identified in the midbrain dataset using the DESeq2 statistical framework.** DEGs with p-value < 0.05 and log2 fold-change > 1.00 are shown.
*See Supplemental_Table3.xlsx*
